# Effects of relocation on Immunological and physiological measures in squirrel monkeys (*Saimiri boliviensis boliviensis*)

**DOI:** 10.1101/2020.10.02.323360

**Authors:** Pramod Nehete, Bharti P Nehete, Greg K Wilkerson, Steve J Schapiro, Lawrence E Williams

**Affiliations:** Department of Comparative Medicine, the University of Texas MD Anderson Cancer Center, Bastrop, Texas, USA; The University of Texas Graduate School of Biomedical Sciences, Houston, Texas, USA; Department of Experimental Medicine, University of Copenhagen, Copenhagen, Denmark

**Keywords:** relocation, stress, Immune and physiological measurements

## Abstract

In the present study, we have quantified the effects of transport, relocation and acclimate/adapt to their new surroundings on squirrel monkey. These responses are measured in blood samples obtained from squirrel monkeys, at different time points relative to their relocation from their old home to their new home. A variety of immunological assays are performed on the phenotype and function of peripheral blood mononuclear cells (PBMCs) in a group of squirrel monkeys that were transported by road for approximately 10 hours from one facility to another. Using a panel of human antibodies and a set of standardized human immune assays, we evaluated the phenotype of lymphocyte subsets by flow, mitogen-specific immune responses of PBMCs in vitro, and levels of cytokines at various time points including immediately before transport, immediately upon arrival, and after approximately 150 days of acclimation. We observed significant changes in T cells and subsets, NK and B cells (CD4^+^, CD8^+^, CD4^+^/CD8^+^, CD16^+^, and CD20^+^). Mitogen specific (*e.g*. PHA, PWM and LPS) proliferation responses, IFN-g by ELISPOT assay, and cytokines (IL-2, IL-4 and VEGF) significant changes were observed. Changes seen in the serum chemistry measurements mostly complement those seen in the hematology data. The specific goal was to empirically assess the effects of relocation stress in squirrel monkeys in terms of changes in the numbers and functions of various leukocyte subsets in the blood and the amount of time require for acclimating to their new environment. Such data will help to determine when newly arrived animals become available for use in research studies.

## Introduction

The number of nonhuman primates (NHP) used in U.S. biomedical research reached an all-time high in 2018 year, according to data released in late September by the U.S. Department of Agriculture (USDA) (NIH report released in September,2018. Nonhuman Primate Evaluation and Analysis Part 1: Analysis of Future Demand and Supply, September 21, 2018). The rising demand for NHP appears to be driven by researchers studying HIV/AIDS, cancer, the brain, Alzheimer’s disease, addiction, Parkinson’s, obesity/diabetes, and emerging infectious diseases like Zika and Ebola and to learn better ways to prevent negative pregnancy outcomes, including miscarriage, stillbirth, and premature birth. This research is also helping scientists to uncover information that makes human organ transplants easier and more accessible, literally giving new life to those whose kidneys, hearts, and lungs are failing. Squirrel monkeys, small New World NHP, have served an important role in studying pathogenesis of human disease conditions such as Alzheimer’s disease [1–3], malaria [4–6], HIV [7], Creutzfeldt-Jakob disease [8, 9] and Giardia infection [10].

Many of these nonhuman primates are raised at one facility and subsequently transported/relocated to another facility for research purposes. Relocating captive nonhuman primates from a familiar home cage or colony room to a novel environment is a potent psychosocial stressor [11–15]. The new and unfamiliar environment presents a sudden, uncontrollable, and unpredictable change. Manipulations of the environments of captive nonhuman primates often have welfare consequences to the animals, including behavioral effects, and for certain manipulations, physiological effects as well. The processes of transporting, relocating, and acclimatizing nonhuman primates across facilities represent manipulations that are likely to have welfare, behavioral, and physiological consequences to the relocated animals [16–19]. Our group and a few others have undertaken a series of studies that have attempted to quantify 1) the effects of transport and relocation, and 2) the amount of time that is required for NHPs to acclimate to the new environments and management procedures after relocation; whether the relocation was to the next room or halfway around the world [20, 21]. Previously, we have reported on the effects of relocation and transport on immunological measurements in chimpanzees, rhesus and cynomolgus monkeys [22–24]. During the winter of 2008 two colonies of different species of nonhuman primates, approximately 500 squirrel monkeys (*Saimiri sciureus sciureus*, *S. boliviensis boliviensis*, & *S. boliviensis peruviensis*) were transported to the University of Texas MD Anderson Cancer Center, Michale E. Keeling Center for Comparative Medicine and Research at Bastrop, TX (KCCMR) from the University of South Alabama in Mobile, AL.

In the present study, we assessed the effects of relocation stress on dependent variables of relevance in research (i.e., immune responses and hematological and chemistry values) in squirrel monkeys. We measured the physiological indicators of stress associated with relocation and the time course of adaptation to the physical and social environments of the new setting. The focus of this project was to empirically assess the effects of transport and relocation on physiological responses in NHPs, and to quantify acclimation processes.

Measuring the effects of transport, relocation, and acclimation should allow investigators to conduct studies that are minimally influenced/confounded by such manipulations.

## Materials and Methods

### Animals, Care, Diet and Housing

Subjects were 30 female squirrel monkeys *(Saimiri boliviensis)*, aged 3-9 years of age from the breeding colony at University of Texas MD Anderson Cancer Center, Michale E. Keeling Center for Comparative Medicine and Research at Bastrop, TX (KCCMR). Monkeys were socially housed throughout the study period in social groups in two connecting cages that are 4’ wide x 6’ tall x 14’ long. Animals had *ad libitum* access to New World Primate Diet (Purina #5040) and water. In addition, they were fed either fresh fruit or vegetables daily. Specialty foods, such as seeds, peanuts, raisins, yogurt, cereals, frozen juice cups and peanut butter, were distributed daily to them as enrichment. At no time were the subjects ever food or water deprived. Subjects were also provided with destructible enrichment manipulanda and different travel/perching materials on a rotating basis to promote the occurrence of species-typical behaviors. The monkeys were examined by veterinarians before, during study period and determined to be healthy. Experiments were approved by the Institutional Animal Care and Use Committee of The University of Texas MD Anderson Cancer Center and were carried out according to the principles included in the *Guide for the Care and Use of Laboratory Animals*, the provisions of the Animal Welfare Act, PHS Animal Welfare Policy, and the policies of The University of Texas MD Anderson Cancer Center [25].

Prior to transport the animals were housed in the Primate Research Laboratory at the University of South Alabama, in social groups had been stable for at least one year. The animals were transported in modified Vari-Kennels^®^ that provided perching and bedding appropriate for squirrel monkeys. Each kennel was provided with a hydration system gel pack, fresh vegetable and fruits, and non-human primate food biscuits. The squirrel monkeys were manually caught and placed in the kennels with a social partner between 1600-1700 hours. Entire social groups were moved at the same time. The trip from Mobile, AL to Bastrop, TX was made in a commercial self-contained, USDA approved, trailer that was climate controlled. The trailer left Mobile, AL between 1700-1730 and arrived in Bastrop, TX 0700-0800 the following morning. Beginning at approximately 0800 the kennels were removed from the trailer and placed in front of their new social housing units. Once an inventory was taken and all animals were found to be present and in the correct social grouping, each kennel was opened, and the animals were released into their new housing. At no point were the animals sedated. The entire colony of 500 animals was moved over a five weeks period following the same routine procedures.

### Collection of samples and Peripheral blood mononuclear cells (PBMC)

Ten animals from three of separate shipments, for a total of 30, were sampled for this study. Baseline blood samples were obtained one day prior to the relocation as part of the animals’ “pre-shipment” physical exam. The next blood samples “Day 1” were obtained within the first few hours upon arrival at the Keeling Center as part of the animals’ first “quarantine” physical exam. The final used for this study was collected at “Day150” post-arrival to monitor the animals’ adaptation to their assigned housing conditions at the Keeling Center. Blood samples (3 mL) were collected, 1.5mL in EDTA anti-coagulant tubes and 1.5 mL in coagulation CBC tubes, from the femoral vein at the different time points. The animals were manually restrained for each of the three blood draws, and no sedatives or other chemical restraints were utilized. All blood sample collections occurred in the morning (8-10AM) before the animals were fed. Before the separation of peripheral blood mononuclear cells (PBMC) from the blood samples, plasma was collected and stored immediately at −80°C until analyzed. The PBMC prepared from the blood samples by the standard ficoll-hypaque density-gradient centrifugation were used for various immune assays [23, 26]. All blood samples were processed at the Keeling Center following domestic overnight shipment (for the pre-travel blood collections) or within 2-4 hours of collection at the Keeling Center (for arrival and day 150 post-arrival samples). The PBMCs, freshly prepared from whole blood collected in EDTA tubes, were more than 90% viable as determined by the trypan blue exclusion method. For each immune assay, 10^5^ cells /well were used for various immune assay.

### Complete blood count and blood chemistry analysis

EDTA whole blood samples were analyzed for a complete blood count on Siemens Advia 120 Hematology Analyzer, Tarrytown, NY. The Parameters analyzed included: total WBC, total RBC, hemoglobin, hematocrit, RBC indices, WBC differential counts, and platelet count. Serum Chemistry was analyzed on Chemistry Analyzer (Beckman Coulter AU680^®^ Chemistry Analyzer). The different parameters analyzed were for example, Glucose, Na, K, Cl, CO2, Cholesterol, Triglycerides, Iron, BUN, Creatinine etc.).

### Flow cytometry

A series of commercially available human monoclonal antibodies that cross react with nonhuman primate mononuclear cells were used in flow cytometry analyses as described previously [10, 26–28]. Briefly, 100μl of whole blood from each sample was added to each 12mm×75mm polystyrene tube (Falcon, Lincoln Park, NJ, USA) containing pre-added cocktail of monoclonal antibodies against CD3, CD4 and CD8 CD16 and CD20 (CD3−PE, clone SP-34; CD4-PerCP, clone L200; CD8-FITC, clone SK1, CD16 APC (clone 3G8), and CD20-APC, clone L27 (all from BD Biosciences, San Diego, USA) and incubated for 15 min at room temperature in the dark. Red blood cells were lysed with 1x RBC lysing solution (Becton Dickinson, USA) following the manufacturer’s instructions. The samples were washed thoroughly in 1x phosphate-buffer saline (PBS) by centrifugation; then cell sediments were suspended in 1% paraformaldehyde buffer (300ul) and acquired on a on a Fluorescence Activated Cell Sorter (FACS) Calibur flow cytometer (BD Biosciences, San Jose, CA, USA). All samples acquired in this study were compensated using the single-color stained cells. Lymphocytes that were gated on forward scatter versus side scatter dot plot were used to analyze CD3^+^, CD4^+^, and CD8^+^ T cell and CD20^+^ B cell lymphocyte subsets using FlowJo software (Tree Star, Inc., Ashland, OR, USA).

For analysis of NK cells, a separate tube with 100ul of blood was stained with separate cocktail of consisting of -CD3 PE (clone SP-34 and CD16 APC (clone 3G8), (all from BD Pharmingen, San Jose, USA) antibodies, as described above. The gating scheme for T-cell, B and NK markers in peripheral blood from a representative cynomolgus macaques has been identified previously [10]. The absolute number of lymphocytes and monocytes, as obtained from hematologic analysis, was used to convert the percentages identified through FACS analysis into absolute numbers for each of the lymphocyte and monocyte subset populations.

### *in vitro* mitogen stimulation of PBMC

The PBMCs, freshly prepared from whole blood collected in EDTA tubes, were more than 90% viable as determined by the trypan blue exclusion method. For each immune assay, 10^5^ cells /well were used for various immune assay. Briefly, aliquots of PBMCs (10^5^/well) were seeded in triplicate wells of 96-well, flat-bottom plates and individually stimulated with the mitogens phytohemagglutinin (PHA), lipopolysaccharide (LPS), and pokeweed mitogen (PWM) (Sigma, St Louis, MO, USA), each at 1 μg/mL final concentration. The culture medium without added mitogens served as a negative control.

### Proliferation Assay

The proliferation of PBMC samples from the monkeys obtained at different time points during the study were determined by the standard [^3^H] thymidine incorporation assay, using mitogens PHA, LPS and PWM (each at 1 ug/mL final concentration). The culture medium served as negative control. Aliquots of the PBMC (10^5^/well) were suspended in RPMI-1640 culture medium supplemented with 10% fetal calf serum and seeded in triplicate wells of U-bottom 96-well plates and incubated with mitogens for 72hr at 37°C in humidified 5% CO_2_ atmosphere. During the last 16-18 hr., 1 μCr of ^3^H thymidine was added. Cells were harvested onto filter strips for estimating ^3^H-incoporation and counted using a liquid scintillation counter. The proliferative response in terms of stimulation index (SI) was calculated as fold-increase in the radioactivity over that of the cells cultured in medium alone. The responses to antigens were considered positive when the SI values were ≤ 2.0 [29, 30].

### ELISPOT Assay for Detecting IFN-γ producing Cells

Freshly-prepared PBMC were stimulated with the different mitogens (PHA, PWM and LPS) to determine the numbers of IFN-γ producing cells by the ELISPOT assay using the methodology reported earlier [10, 31, 32]. Briefly, aliquots of PBMC (10^5^/well) were seeded in duplicate wells of 96-well plates (MABTECH) pre-coated with the primary IFN-γ antibody and stimulated with mitogens PHA, LPS and PWM (each at 1 ug/mL final concentration).

After incubation for 24 hr. at 37°C, the cells were removed, and the wells were thoroughly washed with PBS. Subsequently, 100 uL of biotinylated secondary antibody to IFN-γ (detection antibody) was added to the wells for 3 hr. at 37°C followed by avidin-peroxidase treatment for another 30 min. Purple colored spots representing individual cells secreting IFNγ were developed using freshly-prepared substrate (0.3 mg/mL of 3-amino-9-ethyl-carbazole) in 0.1 M sodium acetate buffer, containing 0.015% hydrogen peroxide. Plates were washed to stop color development, and spots were counted by an independent agency (Zellnet Consulting, New Jersey, NJ) using the KS-ELISPOT automatic system (Carl Zeiss, Inc. Thornwood, NY) for the quantitative analysis of the number of IFN-γ spot forming cells (SFC) for 10^5^ input PBMC. Responses were considered positive when the numbers of spot forming cells (SFC) with the test antigen were at least five and were five above the background control values from cells cultured in the medium alone.

### ELISA Assay for Detecting IFN-α producing Cells

Commercial Cytokines kits were used to measure the concentration of IFN-α (PBL Biomedical Laboratories, Piscataway, NJ) in cell culture supernatants following PHA, LPS and PWM (each at 1 ug/mL final concentration) stimulation of PBMC from squirrel monkeys. ELISA for IFN-α cytokine were performed according to the manufacturer’s instructions.

The minimum IFN-a detectable concentration was 2.9 pg/mL.

### NK Assay

The natural killer activity (NK) was measured as previously described [33]. Briefly, PBMCs from blood were purified by centrifugation on a Ficoll-Hypaque density gradient as described above. Serial two-fold dilutions of the PBMC (effectors) were mixed with ^51^Cr-labeled target cells K562 in triplicate wells of microtiter plates to attain the E: T ratio of 100:1, 50:1, 25:1 and 12.5:1. After 4-h incubation, 100 μl of supernatant was collected from each well and the amount of ^51^Cr released was determined using the γ-counter. To account for the maximum release, the cells were incubated with 5% Triton X-100. Spontaneous release was determined from target cells incubated without added effector cells. The % of specific lysis was calculated by the following formula:

% Specific lysis = (experimental release-spontaneous release) / (maximum release-spontaneous release) × 100.

### Cytokine multiplex assays

Cytokine were measured in cell-free PBMC supernatant using MILLIPLEX-MAP human cytokine/chemokine magnetic bead panel (EMD Millipore Corporation, Billerica, MA, USA) according to the manufacturer’s instructions. There is 91.4%–98.1% homology between the nucleotide sequences of SQM cytokine genes and published sequences of equivalent human and nonhuman primate genes [34, 35]. Briefly, aliquots of PBMC (10^5^/well) were seeded in duplicate wells of 96-well plates and stimulated for 24 hrs. with mitogens PHA, PWM and LPs (each at 1 ug/mL final concentration) supernatant samples were centrifuged (14,000x g for 5 min) and 25 mL of aliquots were used in assay. The 96-well filter plate was blocked with assay buffer for 10 min at room temperature, washed, and 25 mL of standard or control samples were dispersed into appropriate wells. After adding 25 mL of beads to each well, the plate was incubated on a shaker overnight at 40C. The next day, after washing two times with wash buffer, the plate was incubated with detection antibody for 1 h at room temperature and again incubated with 25 mL of Streptavidin-Phycoerythin for 30 min at room temperature. After washing two times with wash buffer, 150 mL of sheath fluid was added into each well and multianalyte profiling was performed on the Bio-Plex 200 system (Luminex X MAP technology). Calibration microspheres for classification and reporter readings as well as sheath fluid, assay, and wash buffer were also purchased from Bio-Rad (Hercules, CA, USA). Acquired fluorescence data were analyzed by the Bio-Plex manager 5.0 (Bio-Rad, Hercules, CA, USA). All steps of incubations were performed on a shaker. The minimum detectable concentration was calculated by the Multiplex Analyst immunoassay analysis Software from Millipore. Cytokines were measured using Nonhuman Primate Cytokine kit with IFN-γ, IFN-a, IL-1b, IL-2, IL-4, IL-6, MIP-1b, TNF-a, and VEGF, from Millipore Corporation (Billerica, MA) using the cytokine bead array (CBA) methodology according to the manufacturers’ protocols and as described previously [36].

### Statistical comparisons

The CBC, chemistry, and immunological data are analyzed using a series of within-subjects One-way Analyses of Variance. The primary comparisons are across the levels of the independent variable; transport and relocation (pre-transport, immediately after transport and after a 150day acclimation referred to as Pre, Day 1, day 150 samples in the results). Two-tailed tests, appropriate correction factors, and planned comparison techniques are utilized to fully explore the data.

## Results

To understand the effect of transport and relocation on immune responses of PBMCs of squirrel monkeys, we performed detailed analyses of cell-mediated immune responses, including assays for 1) Phenotypic analysis by flow cytometry, 2) proliferation, 3) IFN-γ by ELISPOT and IFN-a by ELISA, in response to stimulation with mitogens (e.g., PHA, PWM, and LPS), 4) cytokines in cell supernatant and 5) complete blood count and serum chemistry analysis before, and after relocation.

The lymphocytes and monocytes were first gated based on forward scatter (FCS) versus side scatter (SSC), and then CD3^+^ T cells, CD14* (monocytes), CD3^−^CD16^+^ (NK) cells, CD3^+^CD16^+^ NKT cells, and CD20^+^ B cells were positively identified. The specificity of staining for the various markers was ascertained according to the isotype control antibody staining used for each pair of combination markers, as shown.

### Influence of relocation on major lymphocyte subsets in the peripheral blood

We first checked cross activity of a large panel of commercially available antibodies from different commercial companies at the concentration recommended by the supplier using blood from squirrel monkeys as described previously [37]. The reactivity was considered positive if the signal obtained gave a dot plot clearly distinct from the negative control dot plot using an isotype-matched antibody. As positive controls, fresh blood obtained from normal healthy rhesus monkey as donors were stained in parallel. Based on the data showing a high degree of cross reactivity of human monoclonal antibodies to different lymphocyte subsets in squirrel monkeys, we began probing into immunological indicators of stress associated with relocation.

Using the monoclonal antibodies listed in Table 1 we determined the levels of the different lymphocyte subsets and established normal value ranges in the blood of a total of 30 adult *Saimiri* monkeys from the breeding unit of MD Anderson Cancer Center at Bastrop. Specifically, we analyzed for the T cells (CD3+), NK cells (CD16+), B cells (CD20+ cells), helper T cells (CD3+, CD4+) and cytotoxic/suppressor T cells (CD3+, CD8+) using human monoclonal antibodies that exhibited cross-reactivity with squirrel monkey PBMC. Details of the specificity, clone names, isotypes and supplier of the commercially available human monoclonal antibodies are shown in Table 1.

**Table 1.** Human specific monoclonal antibodies used for squirrel monkey FACS and its reactivity

| Antibody | Supplier | Clone | Isotype | Reactivity |
| --- | --- | --- | --- | --- |
| CD3 | BD | SP34 | IgG3, $\lambda$ | + |
| CD4 | BD | L200 | IgG1 $\kappa$ | + |
| CD8 | BD | RPA-T8 | IgG1 $\kappa$ | + |
| CD16 | BD | 3G8 | IgG1 $\kappa$ | + |
| CD20 | BD | L27 | IgG1 $\kappa$ | + |

The distribution lymphocyte subsets in blood of squirrel monkeys are shown at day pre shipment, post day 2 and post day150 after relocation in Fig 1B. CD3+ T cell count (F (1.7,25.9) =15, p<0.05) showed significant changes across time with the Day 1 arrival levels lower than the Pre and day 150 post-arrival samples. CD4+ T cell counts (F(1.5,22.6)=105,p<0.05), CD8+ T cell counts (F(1.3,19.3)= 6.54, p<0.05), and CD4+CD8+(double positive) T cell counts (F(1.1,16.6)=226,p<0.05) showed significant changes across time with the day 150 levels significantly higher than both the Pre and Day 1 and Day 150 samples. CD16+ NK cells cell counts (F (1.7,25.6) =5.81, p<0.05) showed significant changes across time with Day 1levels significantly lower than the Pre and Day 1 samples. CD20+ B cells counts (F (1.2,18) =9.58, p<0.05) showed significant changes across time with the Day 1 levels significantly higher than both the Pre and Day 1 samples. CD3+CD16+ NK T cells cell counts (F (1.7,25.3) =5.66, p<0.05) showed significant changes across time with the Day 1 levels significantly lower than Pre.

**Fig 1 (A).**
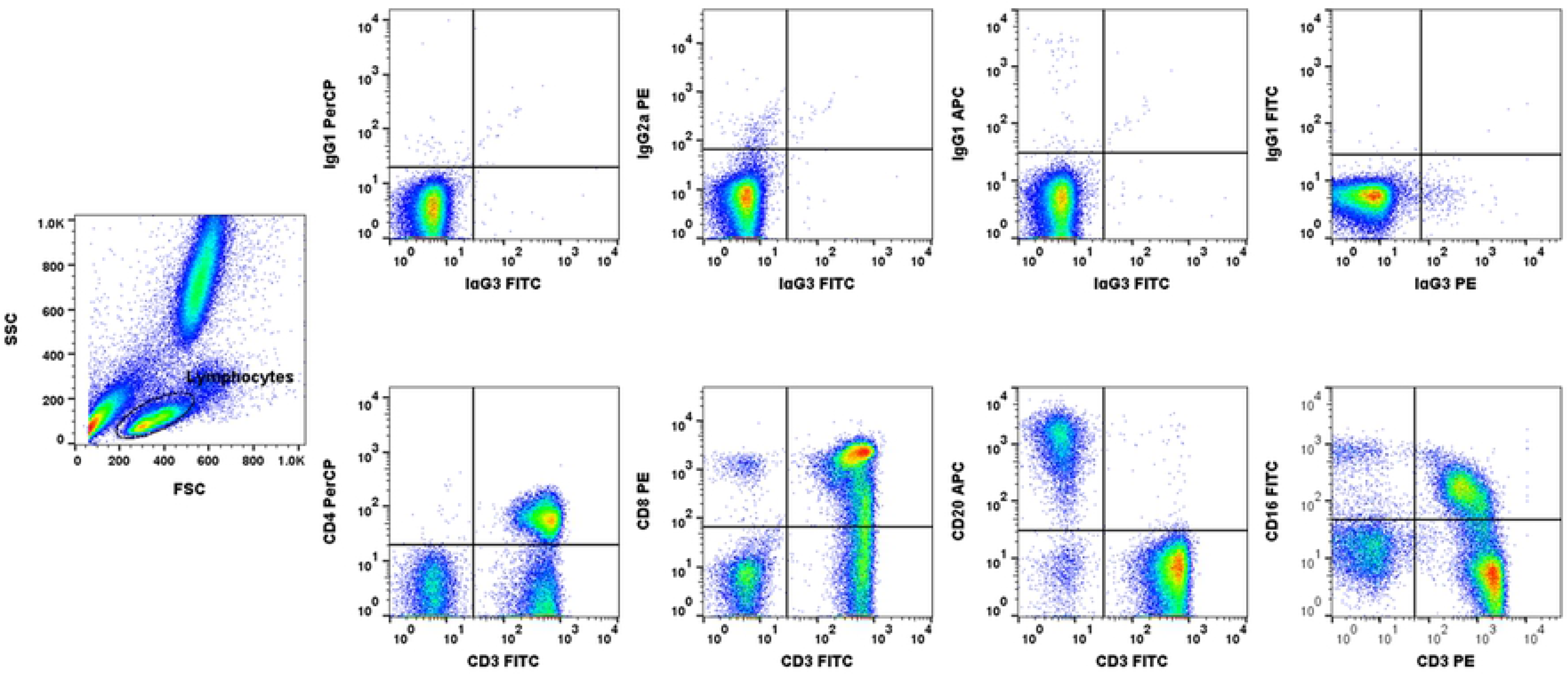
Gating scheme for phenotype analyses of the various cell markers in the peripheral blood from a representative squirrel monkey. The lymphocytes were first gated based on forward scatter (FCS) versus side scatter (SSC), and then CD3^+^, CD4^+^, CD8^+, CD^4+CD8+, and CD20^+^ cells, and NK and NKT cells, were positively identified from the lymphocyte subset. The specificity of staining for the various markers was ascertained according to the isotype control antibody staining used for each pair of combination markers, as shown.

**Figure 1 (B).**
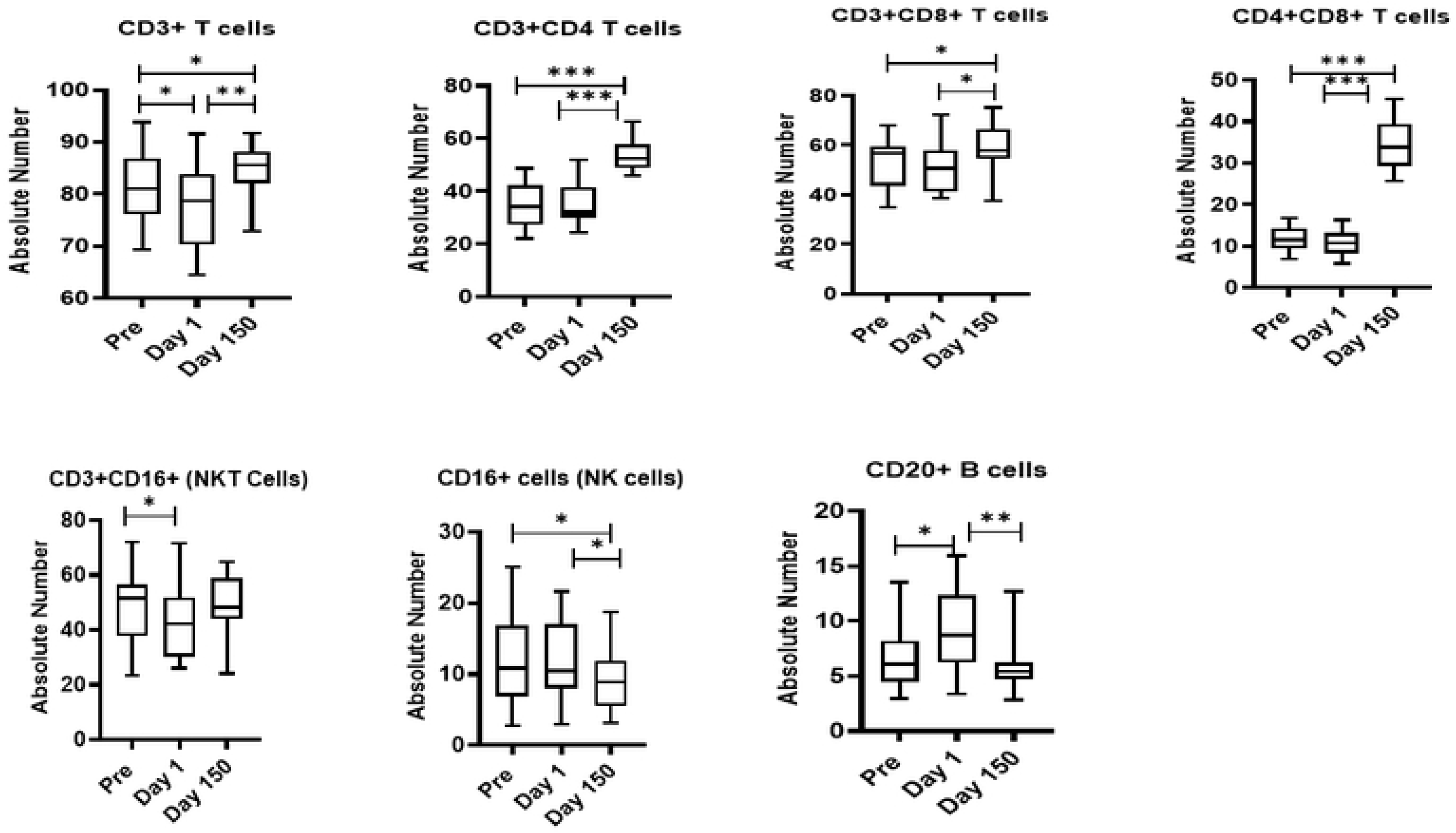
Relocation-dependent differences in lymphocytes in squirrel monkeys. Aliquots of EDTA whole blood were stained with fluorescence-labeled antibodies to the CD3^+^, CD4^+^, CD8^+^, and CD20^+^ to identify lymphocyte subpopulations Pre- and Day 1 and 150 post-transportation and relocation. Values on the Y-axis are absolute lymphocyte cells. P values were considered statistically significant at p<0.05. *Symbol*: * *p<0.05;* \*\**p<0.01;* *** *p<0.0001.*

### Proliferative responses

Since, we found significant differences in expression of T, and B cells, we investigated functional hallmark of proliferation of PBMCs samples from the squirrel monkeys. We measured proliferation in ^3^H thymidine incorporation assay (Fig 2A). The proliferative responses to PHA (F (2,36) =47.1, p<0.0001), and PWM (F (2,51) =33.7, p<0.0001) were significantly higher at post day 2 shipment compare to pre and day150. No statistically significant differences were observed for proliferative response stimulation with LPS (Fig 2A).

**Figure 2 (A).**
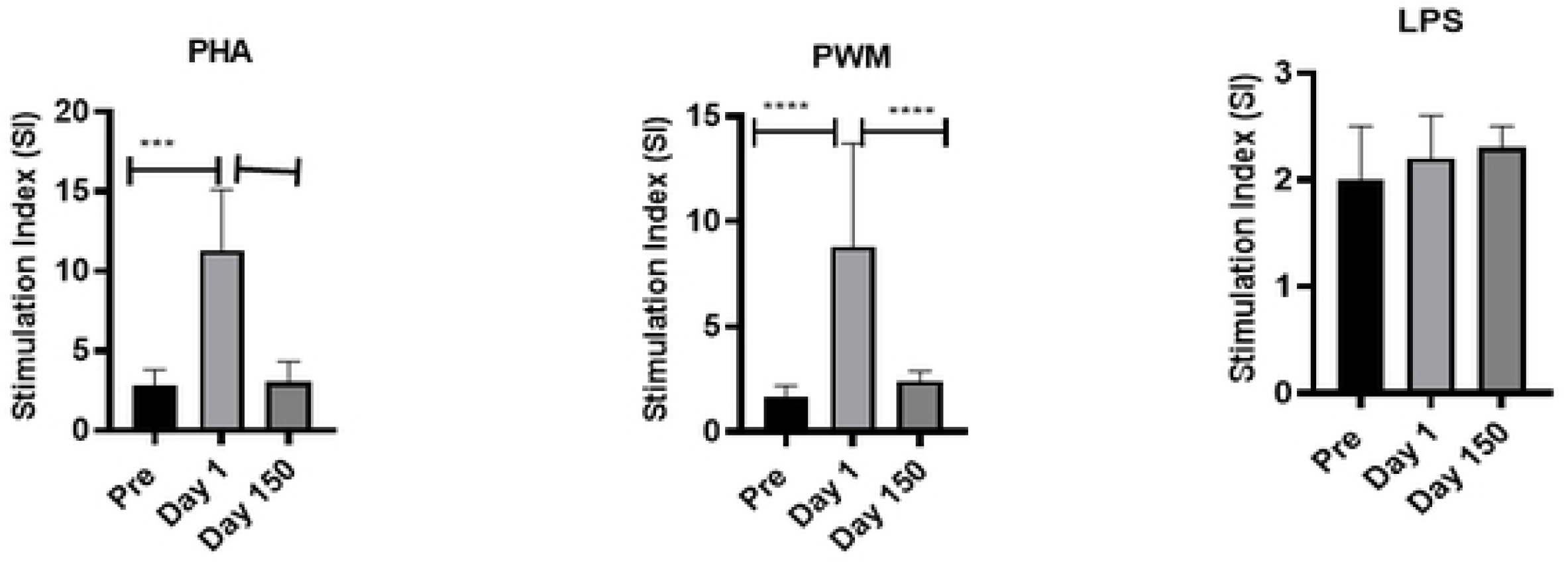
Proliferative response of PBMCs to mitogens. PBMCs that are isolated from blood samples of the squirrel monkeys were used for determining proliferative response to different mitogens, using the standard [3H] thymidine incorporation assay. The proliferation responses are expressed as Stimulation Index (SI) after blank (i.e., medium only) subtraction. P values were considered statistically significant at p<0.05. *Symbol*: * *p<0.05;* \*\**p<0.01;* *** *p<0.0001.*

### ELISPOT assay for detecting mitogen-specific IFN-γ producing cells in squirrel monkeys

Additional functional activity was measured for IFNγ production by PBMCs in response to stimulation with PHA, PWM, and LPS by the cytokine ELISPOT assay. As shown in Fig 2B, squirrel monkey PBMC showed significantly higher numbers of IFN-γ producing cells in response to stimulation with PHA (F(2,51)=136.4,p<0.0001), PWM (F(2,51)=136.4,p<0.0001) and LPS (F(2,51)=113.4,p<0.0001) showed significant changes across time with the Day 150 levels significantly higher than both the Pre and Day 1.

**Figure 2 (B).**
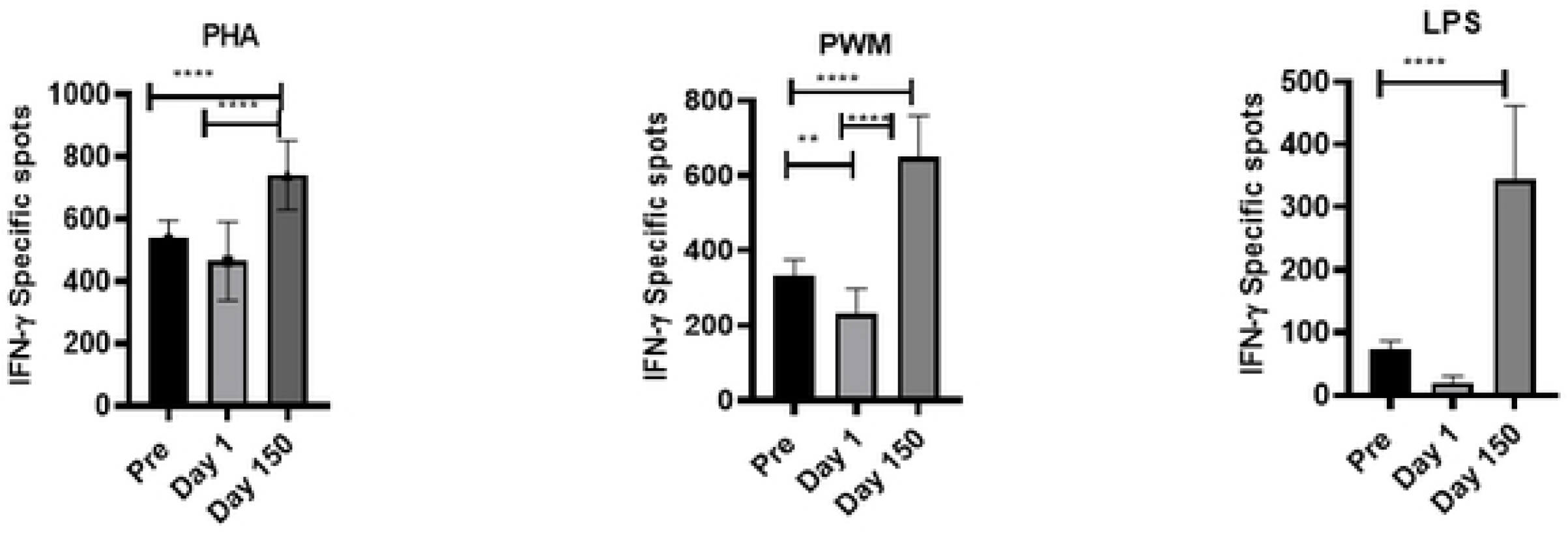
IFN-γ ELISPOT response to mitogens. In duplicate wells of the 96-well microtiter plates, pre-coated with IFN-γ antibody, were seeded with 10^5^ PBMCs from squirrel monkeys stimulated with 1 μg of each of the mitogens for 36 h at 37°C, and then washed and stained with biotinylated second IFN-γ antibody. The total number of spots forming cells (SFCs) in each of the mitogen-stimulated wells was counted and adjusted to control medium as background. See the Methods section for experiment details. P values were considered statistically significant at p<0.05. *Symbol*: * *p<0.05;* \*\**p<0.01;* *** *p<0.0001.*

### ELISA assay for detecting mitogen-specific IFN-α producing cells in squirrel monkeys

Freshly isolated PBMCs were either unstimulated (medium) or stimulated with PHA, PWM and LPS (1*u*g/mL) for 24 hr. and supernatant was collected to measure IFN-α (Fig.3A) by ELISA. In general, Day 150 levels were intermediate between the Pre and Day 1 samples. No significant differences across time were observed for IFN-a ELISA with regard to the PHA, PWM, and LPS assays (Fig 3A).

**Figure 3 (A).**
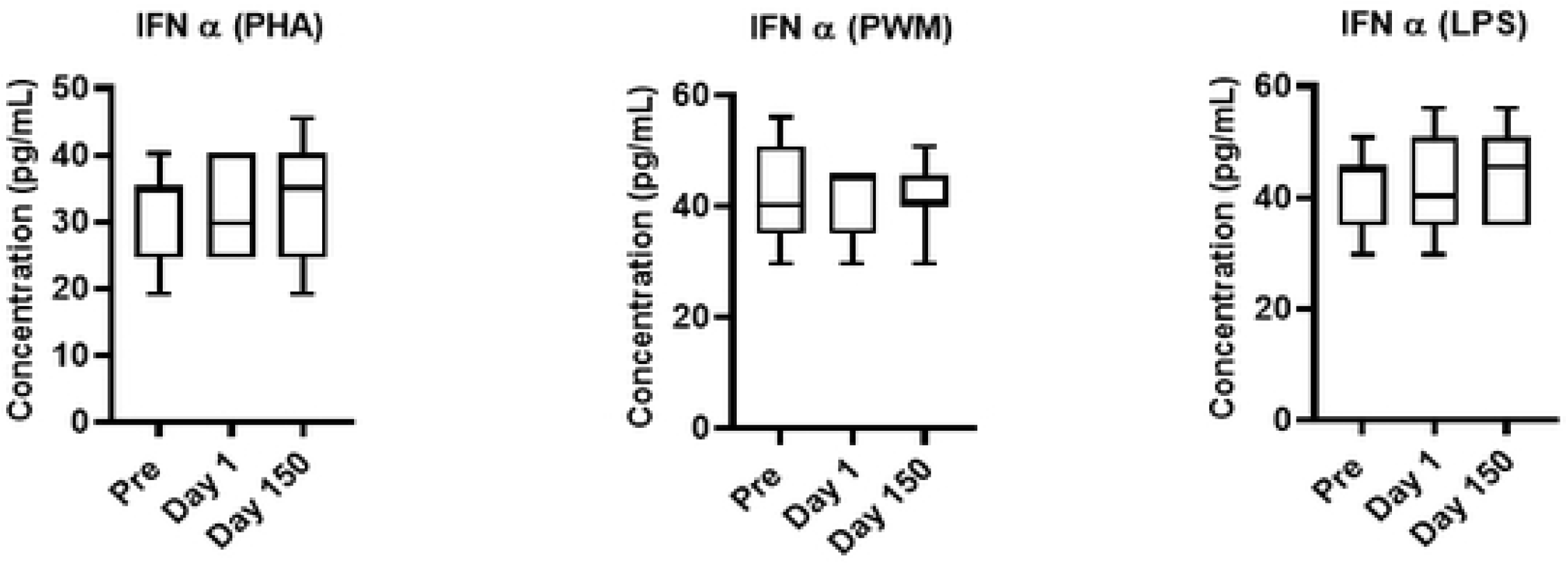
IFN-α **ELISA response to mitogens**. In triplicate wells of the 96-well filter plate, PBMC cultures were stimulated with 1*u*g/mL mitogens for 36hr at 37^°^C and supernatant was collected to measure IFN-α by ELISA. The minimum detectable concentration in pg/mL for IFNα (2.9) was used for considering positive responses. P values were considered statistically significant at *p*<0.05. *Symbol*: * *p<0.05;* \*\**p<0.01;* *** *p<0.0001.*

#### Influence of relocation on natural killer activity

PBMCs from squirrel monkey were analyzed for NK activity using a standard ^51^chromium (Cr) release assay. We observed no significant differences were observed in natural killer activity at different effector to Target ratio (E: T100:1, E: T 50:1, and E: T25:1) (Fig 3B).

**Figure 3 (B).**
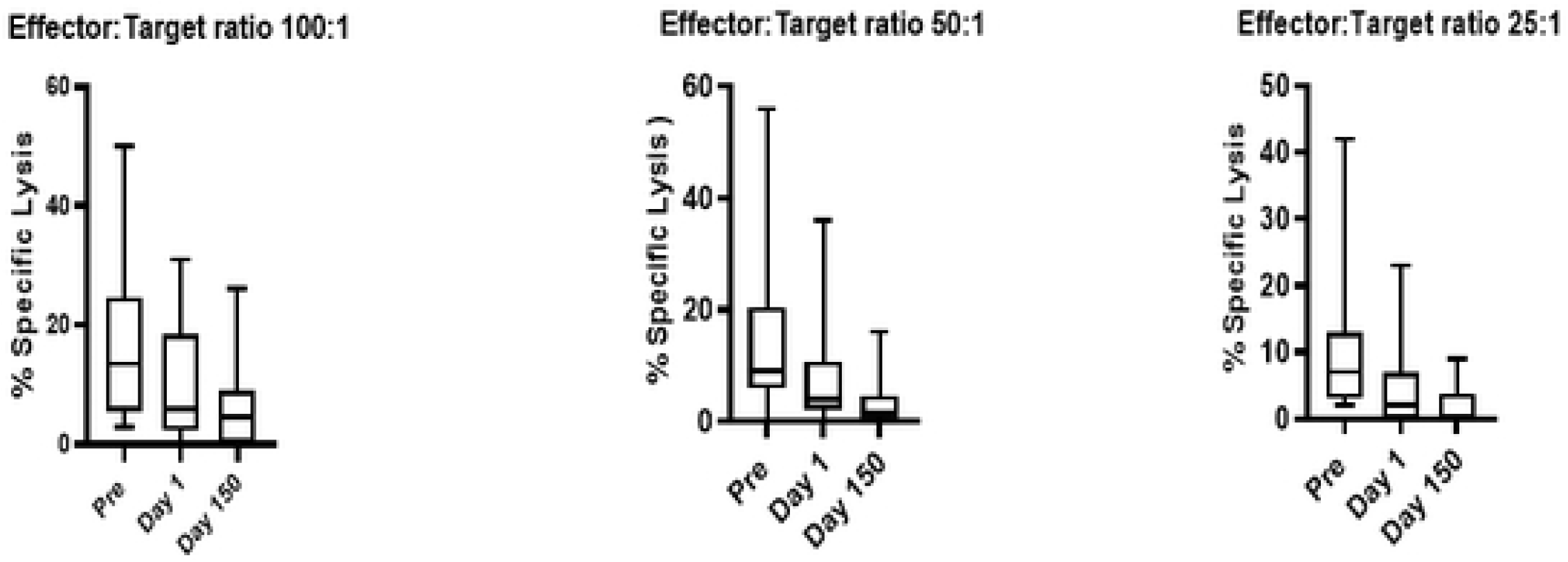
Relocation-dependent differences in natural killer activity. The NK activity of PBMC isolated from squirrel monkeys’ blood was determined by co-culturing with ^51^Cr-labeled K562 target cells at different effector to target cell ratios in culture medium. The percentage (%) of specific lysis is shown at different effector to target ratios. P values p<0.05 were considered statistically significant. *Symbol*: * *p<0.05;* \*\**p<0.01;* *** *p<0.0001.*

### Cytokine multiplex assays

Freshly isolated PBMCs were either unstimulated (medium) or stimulated with PHA, PWM and LPS (1*u*g/mL) for 24 hr. and cell supernatant was collected frozen and used for multiplex cytokines assay using cytokines: IL-1β, IL-2, IL-4, IL-6, MIP-1 β, TNF-α, and VEGF. Only IL-2 (PHA) (F (1.8.25.6) =7.29, p<0.05), IL-4 (PHA)(F (1.8,25.4) =4.3, p<0.05), and VEGF (PHA) (F (1.5,21.6) =5.58, p<0.05) showed significant differences across time with Day 1 arrival levels (Fig 4).

**Figure 4.**
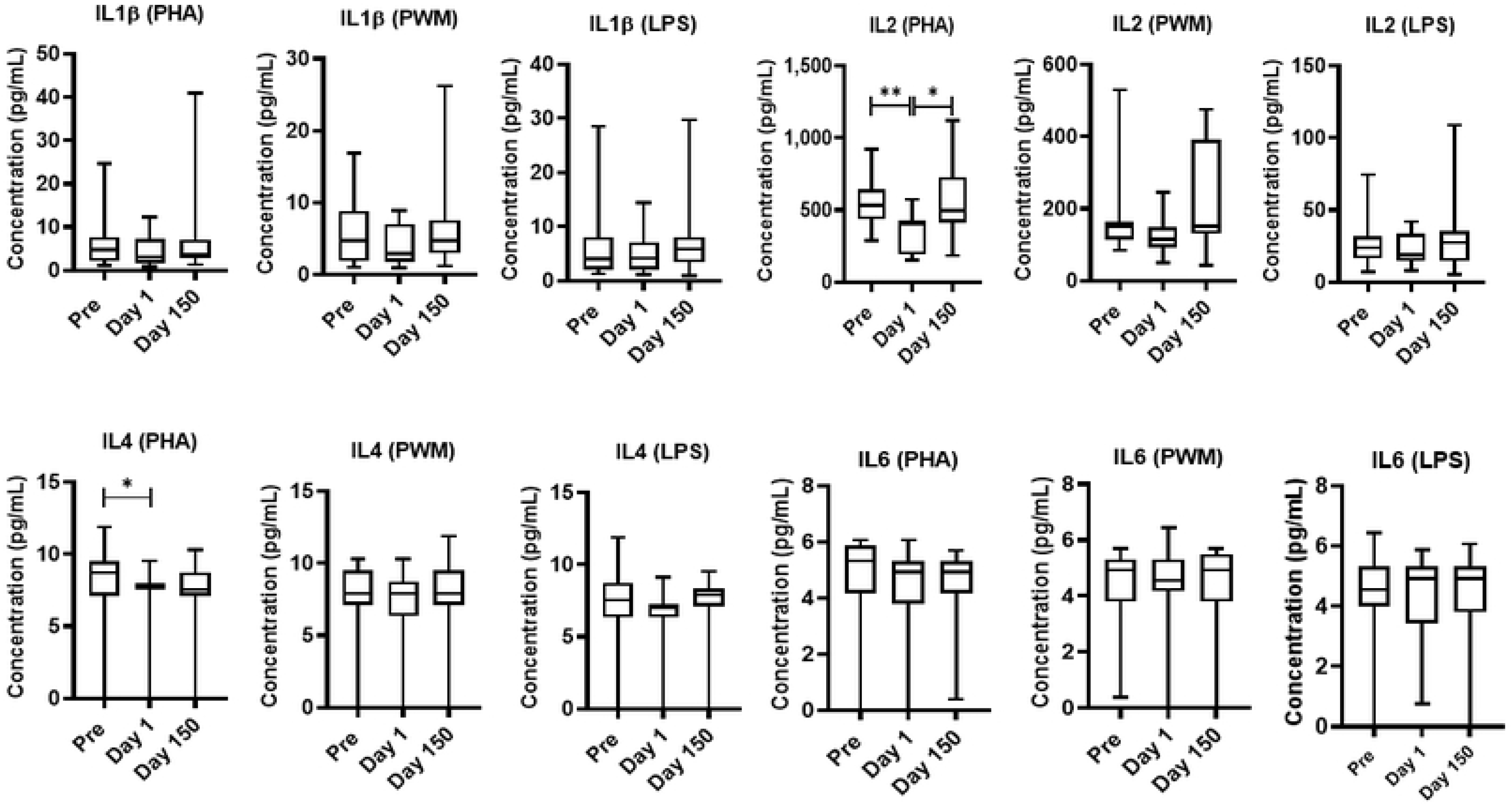
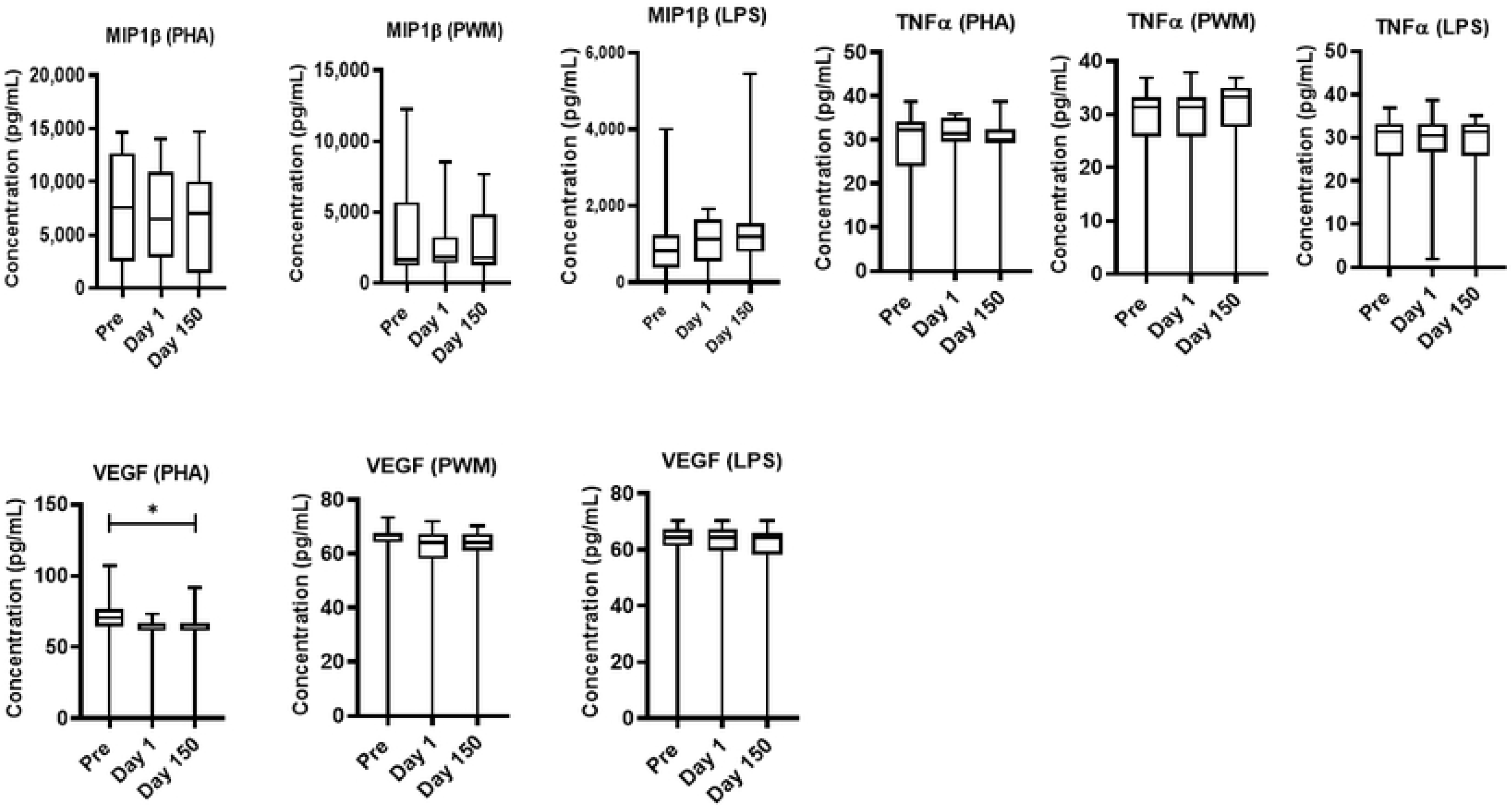
Relocation-dependent differences in Cytokine. In duplicate wells of the 96-well filter plate, 25 μL of cell supernatant was incubated with 25 μL of cytokine coupled beads overnight at 4°C, followed by washing and staining with biotinylated detection antibody. The plates were read on Biorad 200 with use of Luminex technology, and the results are expressed as pg/mL concentration. The minimum detectable concentrations in pg/mL for IL-2 (0.7), IL-4 (2.7), IL-6 (0.3), IL-10 (6.2), IL-12(P40) (1.2), IFN-γ (2.2), and TNF-α (2.1) were used for considering positive responses. See the Methods section for experimental details. P values were considered statistically significant at p<0.05. *Symbol*: * *p<0.05;* \*\**p<0.01;* *** *p<0.0001.*

### Hematology

The blood samples collected at pre shipment Day 1 and Day150 was subjected hematology and analysis is shown in Fig 5A and 5B. The white blood cell count (F (1,12) =12.2, p<0.05) and relative count of lymphocytes (F (1.2,14.4) =16.9, p<0.05) both showed a significant change across time, with the Pre levels significantly higher than both the Day 1 measurements and the post Day 150 samples. The monocyte cell count was significantly different across time (F (1.5,19.5) =5.38, p<0.05) with a significant difference between the Pre levels and Day 150 levels. The neutrophil count (F (1.5,20.3) =50.6, p<0.05) and hematocrit (F (1.8,22.9) =12.6, p<0.05) both showed a significant change across time, with the Pre significantly higher than the post-treatment samples and the Day 150 samples. The eosinophil levels were significantly different across time (F (1.5,19.1) =5.83, p<0.05) with Day 1 levels significantly higher compared to post Day 150 levels. The hemoglobin levels showed significant changes across time (F (1.7,21.3) =9.94, p≤0.05) with Day 1 levels significantly lower than both Pre and Day 1 levels. The red blood cell count (F (1.6,21,1) =8.07, p<0.05), MCHC (F (1.9,24.7) =4.08, p<0.05), and RDW (F (1.7,21.6) =7.59, p<0.05) all were significantly different across time with Pre levels significantly higher than the Day1 levels. MCV, MCH, PLT, MPV, and segmented neutrophil levels showed no significant changes across time.

**Figure 5A-5E.**
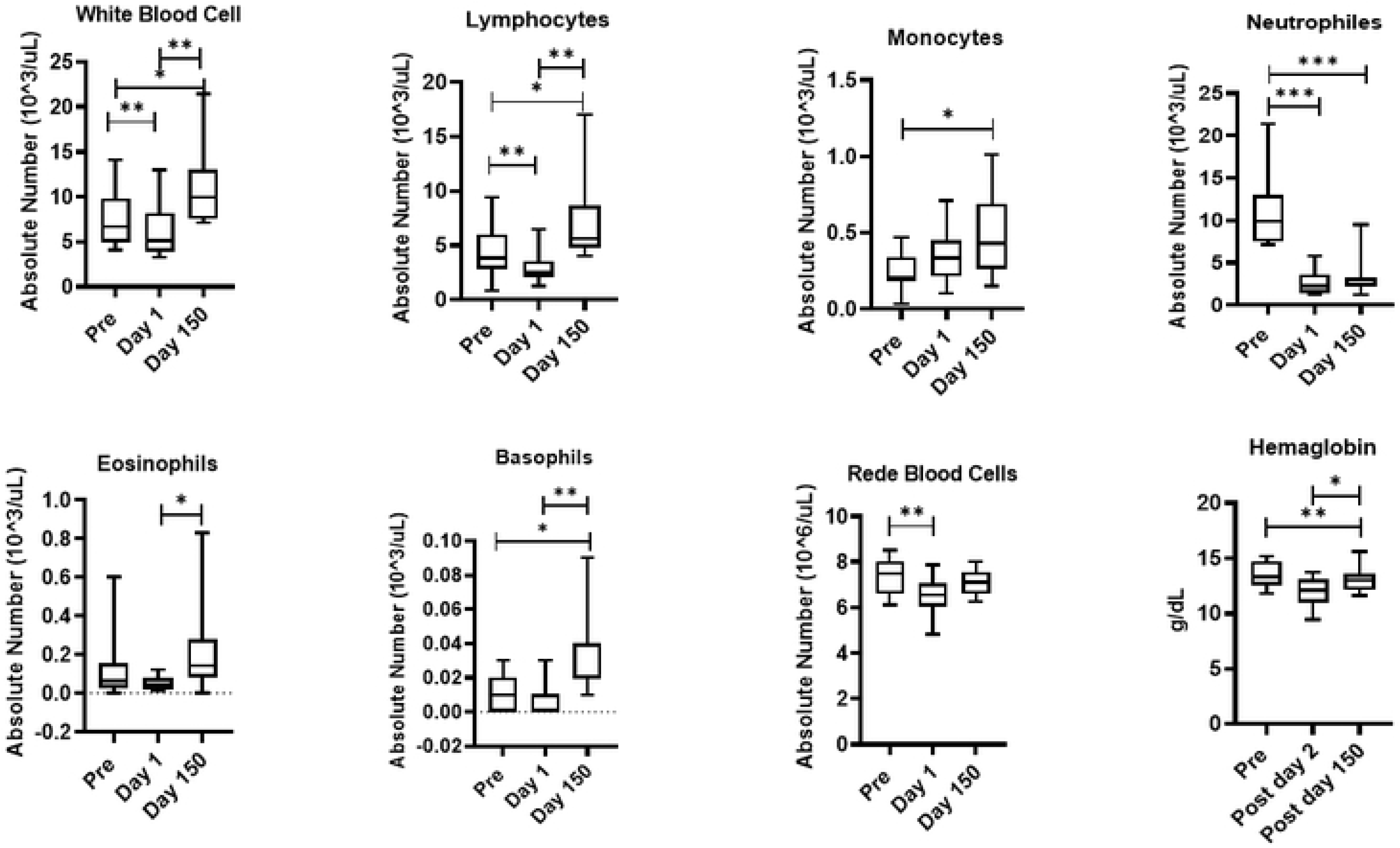
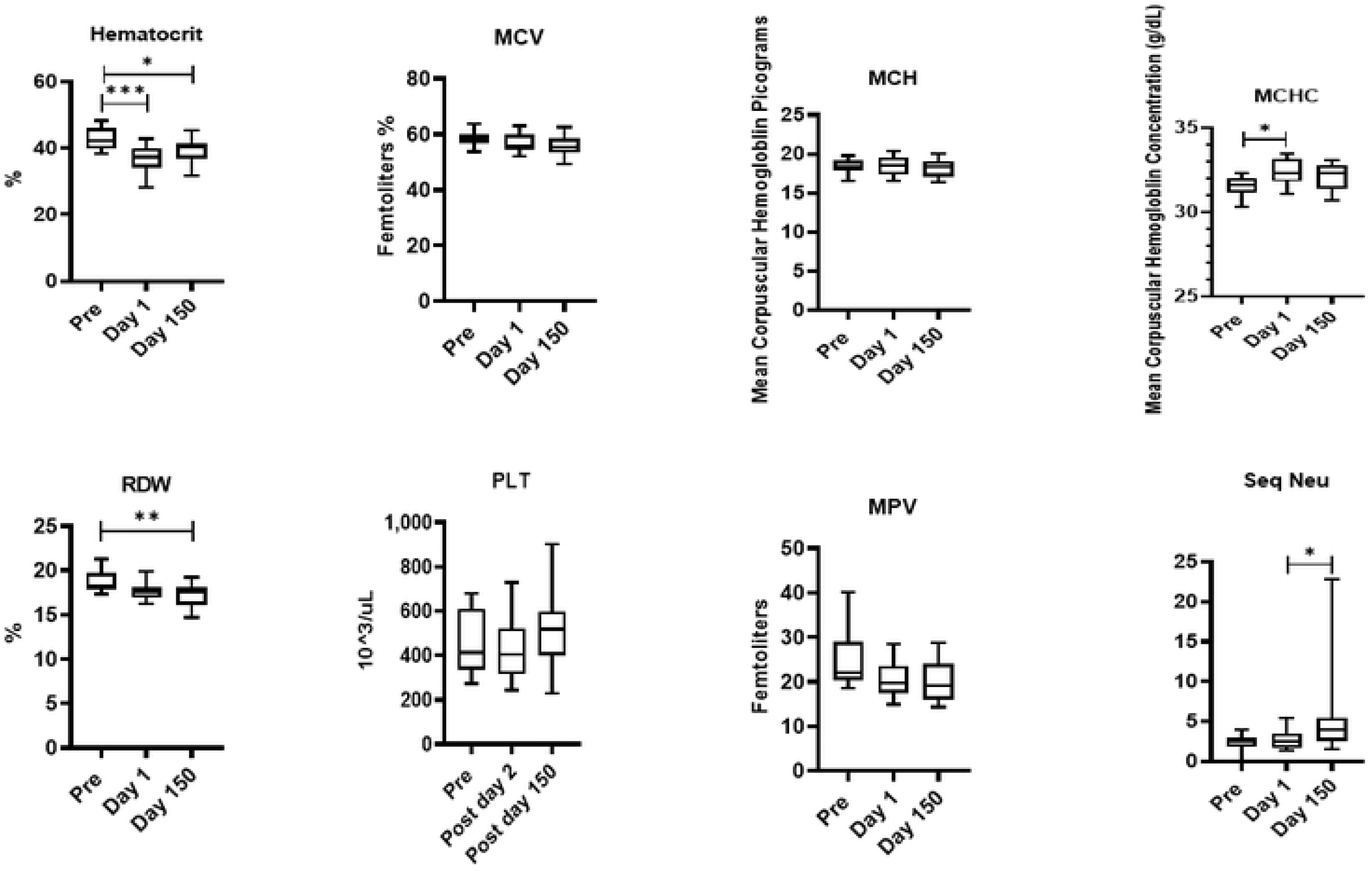
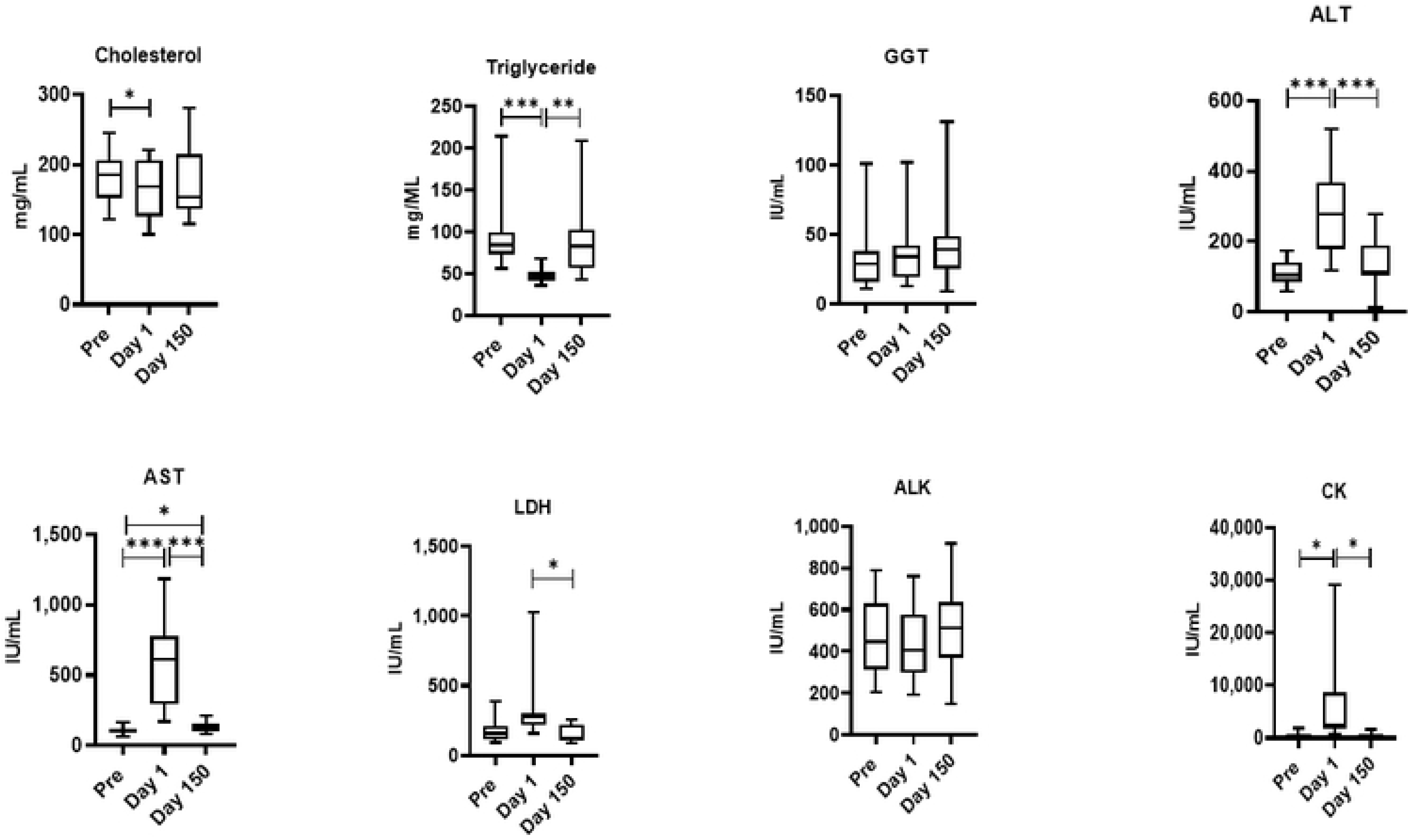
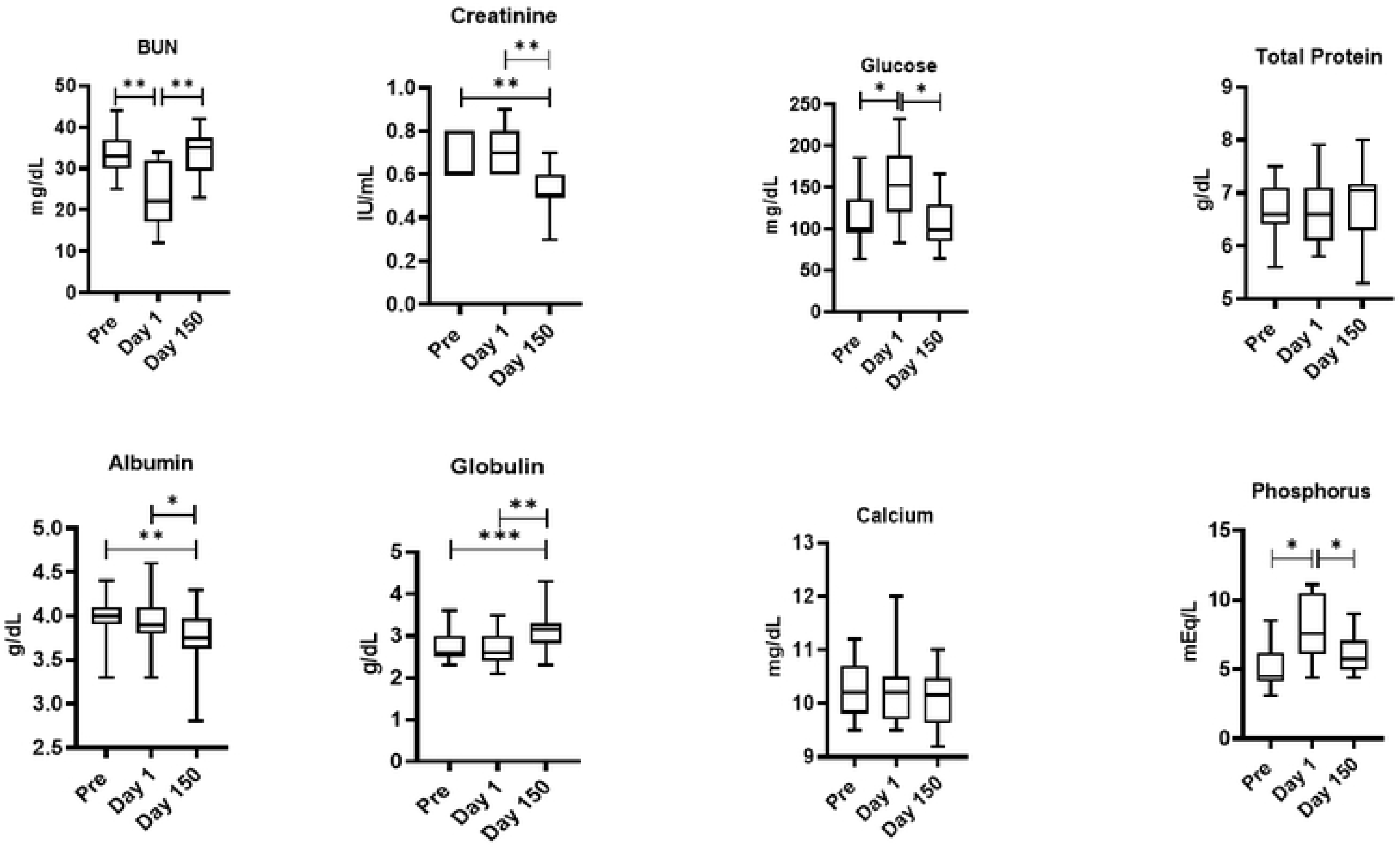
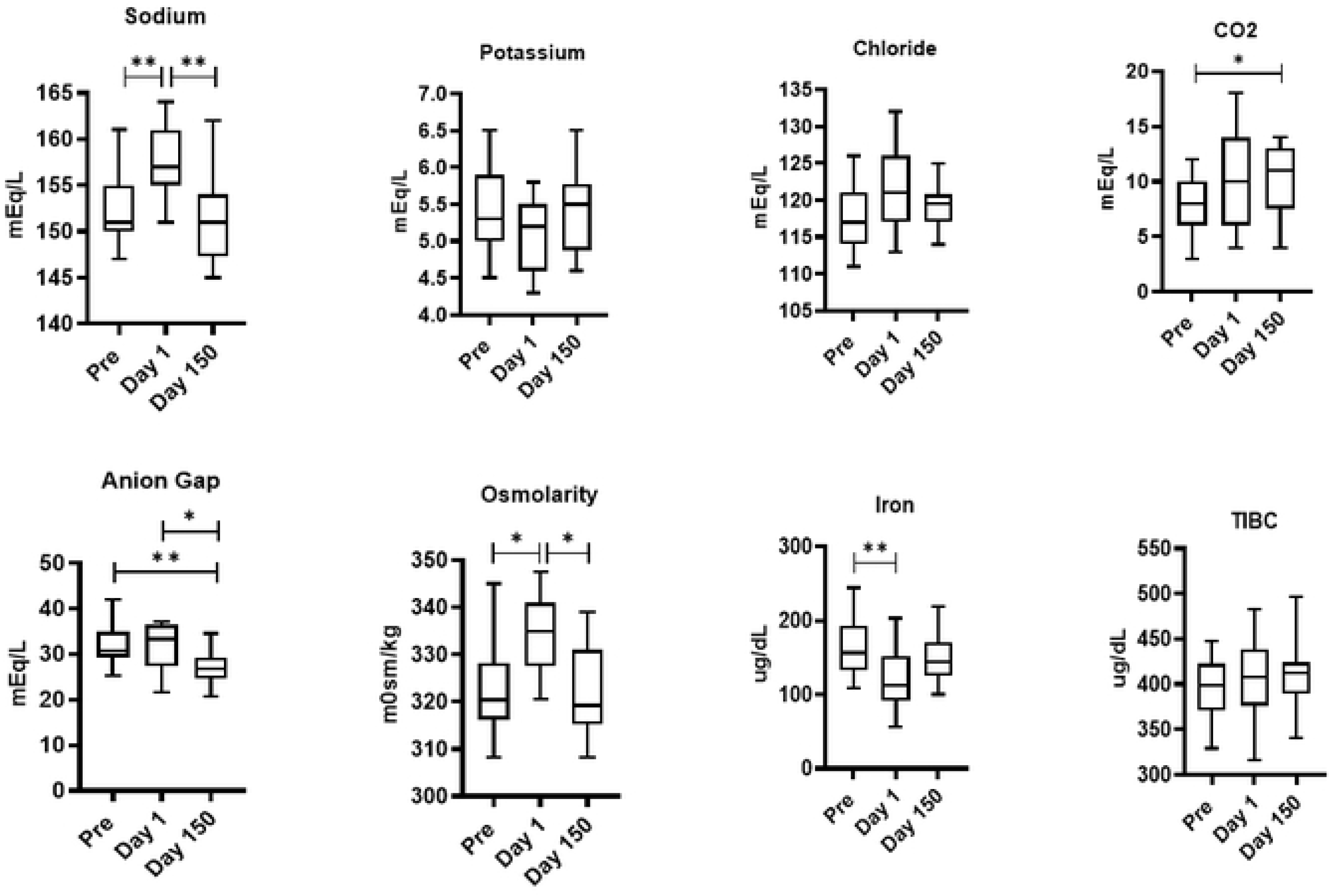
Relocation-dependent differences in Hematology (5A and 5B) and blood Chemistry (5C, D, E). Whole blood by using an automated analyzer Advia (Siemens Healthcare Diagnostics, Tarrytown, NY). Values on the Y-axis are the absolute numbers of lymphocytes and monocytes presented as 10^3 per *u*L of whole blood. P values were considered statistically significant at p<0.05. *Symbol*: * *p<0.05;* \*\**p<0.01;* *** *p<0.0001.*

### Serum Chemistry

Similarly, the blood samples collected at pre shipment, Day 1 and Day 150 was subjected to serum chemistry analysis and is shown in Fig 5C, 5D and 5E. Total bilirubin levels (F(1.3,16.9)=12.8,p<0.05), ALT levels (F(1.7,23.8)=26.7,p<0.05), CK levels (F(1,21.6)=8.9,p<0.05), BUN levels (F(1.4,30.1)=14.6,p<0.05), Glucose levels (F(1.4,19.7)=8.22,p<0.05), Osmolarity (F(1.9,27)=8.09,p<0.05), Phosphorus levels (F(1.4,31.3)=9.96,p<0.05), and Sodium levels (F(1.7,24.5)=12.0,p<0.05) all showed similar significant changes across time, with Day 1 levels significantly higher than Pre and Day 1 and Day 150 samples. Triglyceride levels (F (1.7,24,2) =183, p<0.05) showed a significant change across time with Day 1levels significantly lower than both Pre and the Day 1 and 150 samples. Albumin levels (F (1.6,23.1) =8.58, p<0.05) and Creatinine levels (F (1.6,35.5) =15.2, p<0.05) both showed significant changes across time with Day 1and Day 150 levels significantly lower than both Pre and Day 1 samples. Globulin levels (F (1.5,21.3) =14.6, p<0.05) and Anion Gap (F (1.8,39.4) =5.48, p<0.05) showed similar significant changes across time but with post Day 150 levels significantly higher than both Pre and Day 1 arrival samples. Iron levels (F (1.7,23.9) =5.73, p<0.05) and cholesterol (F (1.2,17.8) =1.07, p<0.05) showed significant changes across time with the Pre samples significantly higher than Day 1 arrival. CO2 levels (F (1.4,31.1) =2.73, p<0.05) showed significant changes across time with post Day 150 levels significantly higher than Pre samples. LDH levels (F (1.8,23.6) =7.44, p<0.05) showed significant changes across time with levels Day 1 arrival levels significantly higher than post Day 150 samples. Pre LDH levels were lower than the Day 1 arrival levels but not significantly so. AST levels (F (1,14.3) =32, p<0.05) showed significant changes across time with Day 1 arrival samples significantly higher than Pre and post Day 150 samples. The post Day 150 AST sample results were also significantly higher than Pre levels. GGT, ALK, Total protein, calcium, potassium, chloride, and TIBC levels showed no significant changes across time.

### Effect of Acclimatization

For most of the hematology measurements all animals had returned to Pre levels at the Day 150 sample. Exceptions include Total Lymphocytes which were higher 150 days post transport (F (1,9) =13.0, ƞ_p_^2^=0.6), and RDW which remained lower than Pre (F (1,9 =7.6, ƞ_p_^2^=0.5).

These hematology effects can mostly be explained by mild dehydration during transport, including changes in the RBC, Hgb, and HCT, even though the animals were provided with water and gel-packs during transport. Transient reductions in the number of WBC’s have been seen with stress as part of intense physical exercise. Unpredictable movements of the truck and crate may have tired and stressed the animals leading to the reduction in WBC’s seen the day of arrival. However, changes in some of the serum chemistry measurements that are related to liver function may be related. Only two hematology measurements showed any post Day 150 effects, higher WBC and lower RDW in the squirrel monkeys. These results are not immediately explainable but do illustrate the types of post Day 150 changes in baseline values that can occur with relocation. Changes seen in the serum chemistry measurements mostly complement those seen in the hematology data. Mild dehydration during the transport may be the causes for changes in Osmolality and sodium levels in the squirrel monkeys. indirectly related may be changes in albumin, phosphorus, and triglycerides that are usually associated with kidney lesions. Elevated levels of CK and LDH, seen are associated with strenuous exercise. Like the fall in WBC’s, this effect is likely due to the animals having to compensate for the unpredictable truck and cage movements. The increase in glucose levels in the squirrel monkeys as well as the decreased levels of iron can be attributed to overall stress and sleep deprivation. Changes in the ALT, AST, and BUN levels in the squirrel monkeys, and total bilirubin in both is usually indicative of liver lesions. Taken together with the platelet changes seen in *squirrel monkeys*, these changes suggest that transport has some transient effect on liver function through a as yet unknown mechanism.

Long term changes seen in BUN, GGT, Chloride, and cholesterol levels in the squirrel monkeys may be an accommodation to the new living arrangements. Although the new cages had a similar volume, they were constructed out of a thermo-neutral material (Tresva^®^) rather than sheet metal. They also received more cage enrichment and were on a different feeding and cleaning schedule. Any of these factors, or other changes in environment and routine, may be enough to alter the physiology of the animals enough to make it different from the original, pre-shipment, baseline.

## Discussion

The data from this study establish clear physiological and immune system effects of transportation and acclimation for squirrel monkeys. Transport involves many factors that can be considered to negatively affect an animal’s physiological state, including anesthesia; loading and unloading; separation from familiar social partners and environments; novel noises, smells, and vibrations; and relocation to a new and unfamiliar environment. The negative effects of these events have been assessed in several species [38–41], as changes in serum levels of cortisol and other physio logical measures, Others have previously studied the effects of transport, relocation, and/or acclimatization on nonhumans primates [11, 24, 41–45], however these studies were conducted on old world and ape species, and focused on limited sets of behavioral or physiological responses and reported a decrease in lymphocytes in cynomolgus macaques, while Kagira, Ngotho (46) found changes in hematological parameters for recently trapped vervet. There are considerably more data available on the effects of transport and relocation in other, non-primate animal species. These reports found similar effects for mice, rats, dogs, and pigs as a function of transport, relocation, and/or acclimatization as suggested by changes in blood glucose, cholesterol, and blood urea nitrogen [47, 48]; lymphocyte counts [38]; and white blood cell counts, body weights, and natural killer cell activity [15, 40].

The data from the present study provide some insight into the time that it takes squirrel to acclimatize to their new surroundings. Some standard clinical chemistry and hematologic values appeared to return to Pre levels by about 6 weeks after arrival, while others did not. Some of the cell-mediated immune responses that were affected by transport and relocation also did not return to Pre levels. The squirrel monkeys were still affected by the transport process weeks after transport and relocation, and probably should not be considered acclimated to their new facility. We reported similar effects in chimpanzees, rhesus and cynomolgus monkey’s relocation to Texas [23, 24]. Their conclusion was that the chimpanzees, rhesus and cynomolgus monkeys should not serve as subjects in studies that use the measured parameters as dependent variables, until they have had adequate time to adjust to their new conditions. This required at least 6-8 weeks for the chimpanzees and longer for the monkeys in this study. The current data does not address the time interval between arrival and 150 days. Additional data is needed to fill in this 5-month gap in the monkey data.

This study focused on the effects of transportation and relocation as assessed by a variety of parameters that are also likely to be dependent measures in biomedical investigations. Significant changes as a function of transport may have few clinical implications in healthy animals, yet they may have numerous research implications. For example, changes in red blood cell counts in investigations of malaria [49] or changes in CD4+ counts in immunodeficiency virus investigations [50]. The better the effects of transport are understood, the better the refinements to management procedures can be. The development of more refined management techniques will result in enhanced welfare for the animals [24, 51–54], enhanced abilities to directly test experimental hypotheses, and may ultimately result in important reductions in the number of primate subjects required to effectively test hypotheses.

Future studies will examine smaller-scale movements of squirrel monkeys within the various housing settings of the KCCMR. We have additional data [24, 55]from reasonably-sized samples of transported and relocated chimpanzees and cynomolgus monkeys that also demonstrate that transport and relocation result in statistically significant changes in a variety of hematological, clinical chemistry, and immunological parameters for these species, although the specific measures affected differ by species.

The data presented here crucial to guide researchers in determining an appropriate acclimation period for their study; however, there are additional factors researchers must consider that can influence how long it takes for acclimation to occur. These factors include the intensity and duration of stress, as well as the species, sex, age, genotype, health status, previous life experience, allometric differences, and even the time of year that acclimation occurs.

## Acknowledgement

We thank Virginia Parks and Bethany Brock for providing blood samples for this study. This study is partly supported by the Cattleman for Cancer Research (PN) and the Squirrel Monkey Breeding and Research Resource (LW) at the Michale E Keeling Center for Comparative Medicine and Research at the MD Anderson Cancer Center.

## Author Contributions

PN and LW designed, interpreted, and analyzed data. PN and BN performed the experiments. GW and SS provided additional interpretation.

